# Impaired activation of the prefrontal executive network during working memory processing in multiple sclerosis

**DOI:** 10.1101/2023.12.22.573051

**Authors:** Chiara Rossi, Diego Vidaurre, Lars Costers, Marie B D’hooghe, Fahimeh Akbarian, Miguel D’haeseleer, Mark Woolrich, Guy Nagels, Jeroen Van Schependom

## Abstract

In multiple sclerosis (MS), working memory (WM) impairment occurs soon after disease onset and significantly affects the patient’s quality of life. Functional imaging research in MS aims to investigate the neurophysiological underpinnings of WM impairment. In this context, we utilized a data-driven technique, the time delay embedded-hidden Markov model (TDE-HMM), to extract spectrally defined functional networks in magnetoencephalographic (MEG) data acquired during a WM visual-verbal n-back task. We observed that two networks show an altered activation in RR-MS patients. First, the activation of an early theta prefrontal network linked to stimulus encoding and attentional control significantly decreased in RR-MS compared to HC. This diminished activation correlated with reduced accuracy in task performance in the MS group, suggesting an impaired encoding and learning process. Secondly, a frontoparietal network characterized by beta coupling is activated between 300 and 600 ms after stimulus onset; this resembles the characteristic event-related P300, a cognitive marker extensively explored in EEG studies. The activation of this network is amplified in patients treated with benzodiazepine, in line with the well-known benzodiazepine-induced beta enhancement. Altogether, the TDE-HMM technique extracted task-relevant functional networks showing disease-specific and treatment-related alterations, revealing potential new markers to assess and track WM impairment in MS.

**Highlights:**

- We decomposed the brain dynamics underlying a WM n-back task in data-driven, spectrally defined whole-brain networks in both healthy controls and people with relapsing-remitting-MS (pwMS).
- PwMS showed a significantly decreased activation of an early theta prefrontal network linked to stimulus encoding and attentional control.
- The weaker activation of this prefrontal theta network is correlated with worse task performance.
- A beta frontoparietal network with a P300-like temporal evolution was significantly modulated by the use of benzodiazepines.
- The model distinguished disease-induced and treatment-induced dynamic network alterations.

## 1. Introduction

Multiple sclerosis (MS) is the most common chronic neuroinflammatory disease of the central nervous system (CNS) in young adults.^1^ Demyelination and neurodegeneration of the CNS characterize the complex pathophysiology of this disease, resulting in a vast range of physical, neuropsychiatric, and cognitive symptoms.^1,2^ Evidence of cognitive impairment (CI) appears in about 50% of the people with MS (pwMS), with cognitive deficits predominantly in attention, working memory, or information processing speed.^2–5^ Working memory (WM) consists of encoding, maintaining, and retrieving a limited quantity of information for few seconds or less.^6^ Since these fundamental processes support all cognitive abilities, from language comprehension to mathematical reasoning, WM impairment majorly impacts the patients’ daily activities.^2,3,5^

WM processes are performed via the synergetic activation of prefrontal, parietal, and temporal regions, forming large-scale brain networks that transiently activate throughout the task.^7,8^ Functional magnetic resonance imaging (fMRI) studies have described the spatial configuration of these dynamic networks and their altered activation in pwMS.^5,9,10^ However, the slow hemodynamic response captured by the fMRI signal overlooks the millisecond temporal evolution of the electrophysiological activity underlying cognitive processing.^11^ Additionally, recent studies have observed reduced fMRI signal quality in pwMS, suggesting that MS-induced hypoperfusion may alter the cerebrovascular coupling.^12^

Electrophysiological measurements, instead, sample brain activity with milliseconds temporal resolution and acquire the electric (via electroencephalography, EEG) or magnetic (via magnetoencephalography, MEG) fields resulting directly from the neuronal activity. Mainly studied in EEG data, the WM event-related response shows a positive peak 300 ms after stimulus onset (in the central frontal-parietal line) – the P300.^13^ This peak is delayed and reduced in amplitude in pwMS, making the P300 a recognized cognitive marker for WM impairment in MS.^14–16^ Additionally, MEG time-frequency studies report reduced hippocampal theta and occipital alpha activity associated with worse task performance.^16,17^ This time-frequency description of MS-induced WM alterations focuses on region-specific activity. However, theta and alpha rhythms are known to lead to neural oscillatory synchronizations forming large-scale brain networks.^18^

The more recent MEG literature on MS presents a few dynamic functional connectivity studies, revealing the frequencies and mechanisms by which the transient network connections arise and how the MS-induced alterations are related to behavioural outcomes, i.e. fatigue.^19–21^ However, these studies are mostly based on resting-state (RS) data.^19,20^ Whereas the role of RS networks in cognition has long been established^22^, RS data may lack information over the temporal dynamics of domain-specific cognitive processes. Task data, on the other hand, allows us to link a functional network to WM-specific processes.^23^ In our previous exploratory analysis^24^, we decomposed the WM brain dynamics, recorded by MEG during a visual-verbal n-back task, in transient functional networks that unfold over tens of milliseconds, hence, the temporal resolution of dynamic cognitive processes. While the spatial layout resembles the classical fMRI networks, we also explored the spectral content and event-related activation of each state, unveiling the potential role of each state in the neuropsychological understanding of WM.^24,25^

We hypothesize that, by applying the same methodology on a dataset that also includes task MEG data of pwMS, the TDE-HMM model can help reveal the data-driven functional networks underpinning WM-specific processes, the activation of which is altered in pwMS, leading to a deeper understanding of the MS-induced WM impairment.^23^ In our results, we observe that MS reduces the activation of the early theta prefrontal network associated with stimulus encoding, suggesting that MS affects the early stage of verbal encoding, directly impacting task performance. Additionally, we observe that treatments based on benzodiazepines have a remarkable effect on the network activation, in particular, amplifying the activation of the M300 beta frontoparietal state.

## 2. Methods

### 2.1 Participants

The dataset includes 38 healthy controls (HC) and 70 people with multiple sclerosis (PwMS); the latter were recruited from the national MS centre (Melsbroek, Belgium) and diagnosed with relapsing-remitting MS via the revised McDonalds criteria (2010).^26^ The included pwMS had an expanded disability status scale (EDSS) score equal to or below 6. To assess the effect of benzodiazepines, we split the MS cohort in two: 16 patients with (MS BZD+) and 54 patients without benzodiazepine treatment (MS BZD-), as benzodiazepines significantly alter the brain dynamics.^27^ Table 1 reports the demographics of these two MS cohorts and the healthy controls (HCs) matched by age. The pwMS were excluded if they (a) had experienced a relapse and/or (b) were treated with corticosteroids within six weeks before the start of the study, or (c) if they carried pacemakers or dental wires or (d) suffered from epilepsy or psychiatric disorders.

**Table 1.** Demographics of the dataset. The three groups are HC (healthy controls), MS BZD- (pwMS without BZD treatment), and MS BZD+ (pwMS undergoing BZD treatment). EDSS = expanded disability status scale. To evaluate the difference in age, education level, and neuropsychological scores (for the two tests CVLT-II and SDMT) across the three groups, we used a one-way ANOVA test with treatment status as the grouping variable. To test the difference in disease duration and EDSS score between MS groups, we utilized Wilkinson’s rank sum test. NA = not applicable, NS = not significant.

|  | HC | MS BZD- | MS BZD+ |  |
| --- | --- | --- | --- | --- |
| **Number of subjects**<br>(% male) | 38 (39%) | 54 (47%) | 16 (0%) | NA |
| **Age**<br>mean (SD) | 47.9 (11.8) | 47.5 (9.5) | 47.6 (7) | NS |
| **Education level**<br>mean (SD) | 15 (2) | 14.4 (2.7) | 13 (2.4) | *p <.05<br>(one-way ANOVA test) |
| **Disease duration**<br>median (IQR) | NA | 15 (14) | 15 (10.5) | NS |
| **EDSS**<br>median (IQR) | NA | 2.5 (1.5) | 3.25 (1) | *p <.05<br>(Wilkinson's rank sum test) |
| **California Verbal Learning test II (CVLT-II)**<br>z-score data<br>median (IQR) | 0.182(1.18) | 0.182(1.50) | -0.033(2.09) | NS |
| **Symbol-to-Digit modality test (SDMT)**<br>z-score data<br>median(IQR) | 0.1(1.32) | -0.1(1.5) | -0.28(0.95) | NS |

The patient recruitment started in 2015 and concluded in 2018. All participants signed an informed consent, and the study was approved by the ethical committees of the National MS Center Melsbroek and the University Hospital Brussels (Commissie Medische Ethiek UZ Brussel, B.U.N. 143201423263, 2015/11).

Both pwMS and healthy subjects performed the BICAMS^28^, a battery of neuropsychological tests including the single digit modality test (SDMT) to assess information processing speed, and the California Verbal Learning-II (CVLT). Both tests can be relevant when evaluating working memory functioning during a visual-verbal n-back task. Table 1 reports the neuropsychological scores for the three groups.

### 2.2 Data Acquisition

Participants underwent MRI and MEG acquisitions. The MEG machine was located at the CUB Hôpital Erasme (Brussels, Belgium) in a lightweight magnetically shielded room (MSR, MaxshieldTM, MEGIN Oy, Croton Healthcare, Helsinki, Finland). Because of a system update, 39 subjects were scanned using the Neuromag VectorViewTM system, while 69 were scanned with the NeuromagTM TRIUX system (MEGIN Oy, Croton Healthcare, Helsinki, Finland). The data were acquired using 102 triplets (two planar-gradiometers and one magnetometer) of SQUID sensors. Three coils were attached to the left and right forehead and mastoid to track the head’s movements, and three sensors were used to record electrocardiography (ECG) and electrooculogram (EOG). The subject’s head shape and fiducial (nasion, left and right preauricular) points were registered using an electromagnetic tracker (Fastrak, Polhemus, Colchester, Vermont).

The MRI was acquired with a 3T Achieva scanner (Philips, Best, Netherlands) at the Universitair Ziekenhuis Brussel (Jette, Belgium). The 3D MR images were T1-weighted; the subjects were in a head-first supine position. The scan used an echo pulse sequence gradient with echo sequence TE 2.3030; the recording parameters were TR = 121 4.939 ms, flipping angle 8, field of view 230 × 230 mm2, number of sagittal slices 310 with a 0.53 by 0.53 by 0.5 mm3 resolution (voxel).

The MEG (functional) and MRI (structural) data were collected within a few days (median 5, IQR 2-10 days).

### 2.3 Task data

During the MEG recording, each participant performed a visual-verbal n-back task. During the n-back, a sequence of letters is displayed, and the participants must press a button (right hand) when the target letter appears. A letter identical to the n^th^ preceding one (n = 1,2) becomes a target. If n = 0, the letter X becomes the target. Figure 1 shows a graphic representation of the task. For each load condition (n = 0,1,2), the experiment consisted of four blocks of 20 letters (20% as target stimuli) presented pseudo-randomly. Before the actual recording, each participant underwent a training session. The letters were projected on a 6x6.5 cm screen located 72 cm in front of the subject. A photodiode attached to the screen precisely detected the stimulus onsets.

**Figure 1.**
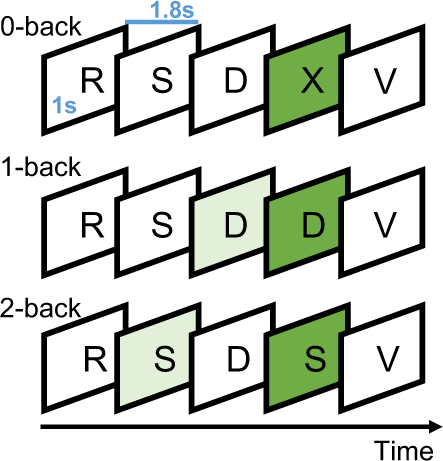
Schematic illustration of the visual-verbal n-back task. The dark green squares highlight the letters shown, which are target trials, and the light green squares represent the letters that are matched with the target to identify the trials. From the top, each line depicts one paradigm condition, 0, 1, and 2 back, respectively. Each letter is shown for 1 second, and the inter-stimuli time is 1.8 seconds.

We only included the correctly identified trials, discarding the missed targets and distractors with an answer. The reaction time is evaluated as the time between the target onset and the moment subjects pressed the button. We also evaluated the accuracy of performance, measured as the ratio between the number of correctly identified trials and the total number of target trials.

### 2.4 MEG data preparation

#### MEG Preprocessing

The raw data was first cleaned for background noise and head movements using the temporal extension of the maxfilter software, bandpass filtered in [0.1 330] Hz (version 2.2 with default parameters; MEGIN Oy, Croton Healthcare, Helsinki, Finland). The preprocessing was implemented in MATLAB 2020b using the OSL package.^29^ The preprocessing pipeline follows ref.^30^ The MEG data were coregistered to the subjects’ T1 MRI via RHINO coregistration using the subject-specific fiducial points traced by the Polhemus electromagnetic tracker. Afterwards, the data were downsampled from 1000 Hz (acquisition rate) to 250 Hz and bandpass filtered to [1, 45] Hz. A notch filter at 50 Hz was introduced to remove the remaining powerline noise that could interfere with the model implemented.

Artefact removal was performed via independent components analysis (ICA), considering 62 components, and the components that correlated (r>0.5) with the ECG or EOG recordings were discarded. Finally, the clean signals and rejected components were manually checked and visually inspected.

The data acquired by different MEG sensors (magnetometers and planar gradiometers) were normalized to overcome the difference in data variance across sensor types, following.^31^ Afterwards, we applied a bilateral beamformer, based on a Bayesian principal components analysis as developed in^31^, to reconstruct the sources of the acquired signals. The source reconstruction was based on a single-shell forward model in the MNI space with a projection on a 5 mm dipole grid.

#### Parcellation

The MEG source-reconstructed data were parcelled using a 42 region-of-interest (ROIs) atlas used before in.^20,24,32^ This data-driven parcellation includes only cortical regions and the first principal component between the voxels’ time series was assigned to the parcel. The ROIs’ time courses were orthogonalized by multivariate symmetric leakage correction to discard signal leakage across parcels.^33^ Lastly, we applied the sign flipping algorithm, as proposed by Vidaurre et al^32^, to overcome the sign ambiguity issue affecting beamformed data.

### 2.5 TDE-HMM

We implemented the time delay embedded-hidden Markov model (TDE-HMM) to extract dynamic functional networks from the MEG preprocessed data. For a thorough mathematical description of the model, we refer to.^25,34^ We have previously implemented the same technique to unveil the network dynamics underlying working memory processing in healthy subjects^24^, so we only describe the method conceptually in what follows.

#### Time delay embedded - Hidden Markov model

Generally, a hidden Markov model describes the empirical data (MEG) as alternating activations of a discrete number of hidden states, the functional brain networks, as illustrated in Figure 2. As a Markovian model, the activation of a state at time t depends directly on the observed data at time t and the adjacent state activated at time t-1.^35^ Here, we use the time delay embedded-HMM, in which each state is modelled by a (low-rank) Gaussian distribution across regions and time points, spanning through a time window of 60 ms (15 time points), thereby capturing specific spectral patterns across regions. The Gaussian distribution is assumed to have the mean equal to zero so that the covariance across regions and time points is the only state parameter. The estimation of the model parameters is done using stochastic Bayesian variational inference.^32,34^

**Figure 2.**
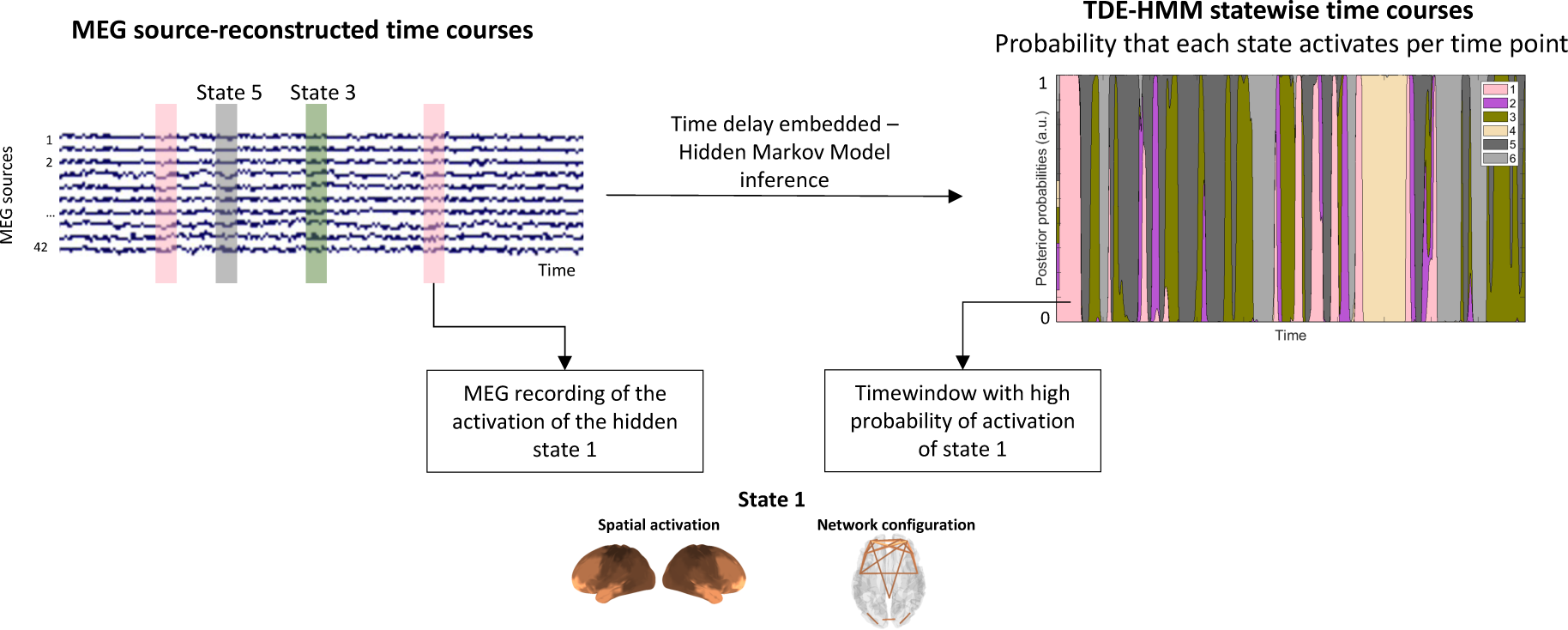
Illustration of the relationship between MEG data and the HMM states. The HMM inference extracts the spectrally defined spatial distribution of power and coherence. The activation of a state results in the MEG recordings, which is translated in statewise time course of activation by the HMM inference, revealing the underlying transient waxing and waning of the hidden states.

We ran the TDE-HMM at the group level, i.e. on both the HC and MS concatenated data, so that the brain states are straightforwardly comparable between groups. Although the states are inferred at the group level, the statewise time, space, and frequency profiles can be evaluated per subject (as explained below), which allows us to compare the states’ features between groups. This is analogous to the dual estimation process.^36^ Regarding the number of states to infer, we set apriori the number to 6, based on previous work.^24^ Importantly, the brain states are inferred without entering any information on the task. The model’s output that we consider is the posterior probability (probability of activation) of each state. This gives us the probability of a state being active at each point of time – forming the states’ time series.

#### Spectral analysis

We extracted the spectral description of each state and subject separately. First, the MEG data were multiplied by the statewise posterior probabilities – resulting in the statewise MEG time courses. From this, a non-parametric multitaper was used to compute the power spectral density (PSD) for each brain region and the coherence (COH) between each pair of brain regions. As the multitaper is run singularly for each subject, we obtain the subject and state-specific spatial (PSD) and phase-coupling (COH) maps.^25,34^

The PSD and coherence were extracted in a broad frequency band [1, 40] Hz. Afterwards, we decomposed the frequency band into four components (spectral mode) in a data-driven way using a non-negative matrix factorization of the coherence measure, as described in.^25,30^ The number of components (4) was chosen considering the number of conventional frequency bands included in the 1-40 Hz spectrum: theta/delta [1-8] Hz, alpha [8-12] Hz, beta [12-25] Hz, and low gamma [25-40] Hz. By multiplying each spectral mode for the PSD and COH of each subject and state, we extracted the spatial (PSD) and phase-coupling (COH) maps for each data-driven spectral mode. To plot, we z-normalised the PSD maps and displayed only the phase-coupling connections that survived thresholding via a Gaussian mixture model (GMM).

### 2.6 Statistical Analysis

#### Event-related ac1va1ons

We epoched the statewise time courses (probability of activation per time point) with respect to the stimulus onset, creating epochs of 1400 ms, [-200, 1200] ms. Each epoch was baseline corrected, considering as baseline the [- 200, -30] ms time window before the stimulus. Next, we ran a generalized linear model (GLM) with 7 regressors: the 6 task conditions (0, 1, 2 back targets and distractors), and a constant regressor. From the results of the constant regressor, we identify the task-relevant states as those that significantly modulate their activation (time course) in response to the task^24^.

Following, we ran a statistical analysis to investigate the group differences (HC vs MS BZD- vs MS BZD+) in the event-related activations for the task-relevant states. We took the average task-evoked response per subject and paradigm condition, considering the 0, 1, and 2 back, separately. We implemented non-parametric statistics, a permutation analysis. In the permutation loop (number of permutations = 1000, a one-way ANOVA test evaluated the difference in task-evoked occupancy level across the three groups (HC, MS BZD-, and MS BZD+). The permutation test was computed per state, per paradigm condition (0, 1, and 2 back), and per time point over the time window [0, 1000] ms. Then, we corrected the tests for multiple comparisons via maximum statistics (maximum over time and paradigm conditions). This analysis provided the time points within an epoch in which, for each state singularly, the event-related activations of the three groups (HC, MS BZD-, and MS BZD+) significantly differ.

#### ER features – amplitude peak and latency

We ran a peak analysis to further explore the morphology of the states’ event-related activations and the differences between groups. We extracted the peaks of states 1 and 5 as these were the only ones to show significant differences between groups in the ER activations. For state 1, the maximum positive peak was extracted in the [100, 300] ms time window, whilst the M300 maximum positive peak of state 5 was extracted in the [300, 600] ms time window. We also extracted the early negative peak in state 5 in the time window [0, 200] ms as it showed a significant modulation between groups in the ER profile. We are interested in observing the amplitude and latency for each peak.

We ran an n-way ANOVA test to investigate whether the peaks’ measures (latencies and amplitude, separately) differ between groups (HC, MS BDZ-, and MS BZD+) or across paradigm conditions (0, 1, 2 back). The n-way ANOVA tests were all corrected for multiple comparisons via false discovery rate. Finally, we utilized Tukey’s HSD (honestly significant difference) test to identify which groups were leading the significant difference.

This peak analysis served to link the findings over the states’ altered ER activations in the MS groups to the observable behavioral data. After extracting the peaks’ amplitude and latencies, we correlated them (Pearson’s correlation) with the behavioral data: mean reaction time, mean accuracy, SDMT, and CVLT tests. The correlation was run per state, peak, and task condition (0,1, 2 back), separately. Given the exploratory nature of this analysis, we did not correct the correlations for multiple comparisons.

## 3. Results

### 3.1 Task Performance

Figure 3 reports the distribution of the subjects’ mean reaction time and accuracy for the three groups (HC, MS BZD-, and MS BZD+) and for each task condition (0,1, and 2 back). The mean reaction time differs significantly between the HC and MS BZD- groups in the 2-back condition (one-way ANOVA test, F = 4.82, p<0.05, multiple comparison correction via FDR, and following Tukey’s HSD Test, p<.05, 95% CI = [-.107, -.0104]), Figure 4a. In the 1-back condition, instead, the accuracy significantly differs between groups (one-way ANOVA test, F = 10.76, p<.05, multiple comparison correction via FDR), Figure 4b. In particular, the mean accuracy of the MS BZD+ group is significantly different from both the mean accuracy of the HC group (p<.05, 95% CI = [.057, .19]) and the MS BDZ- group (p<.05, 95% CI = [-.05, -.18]).

**Figure 3.**
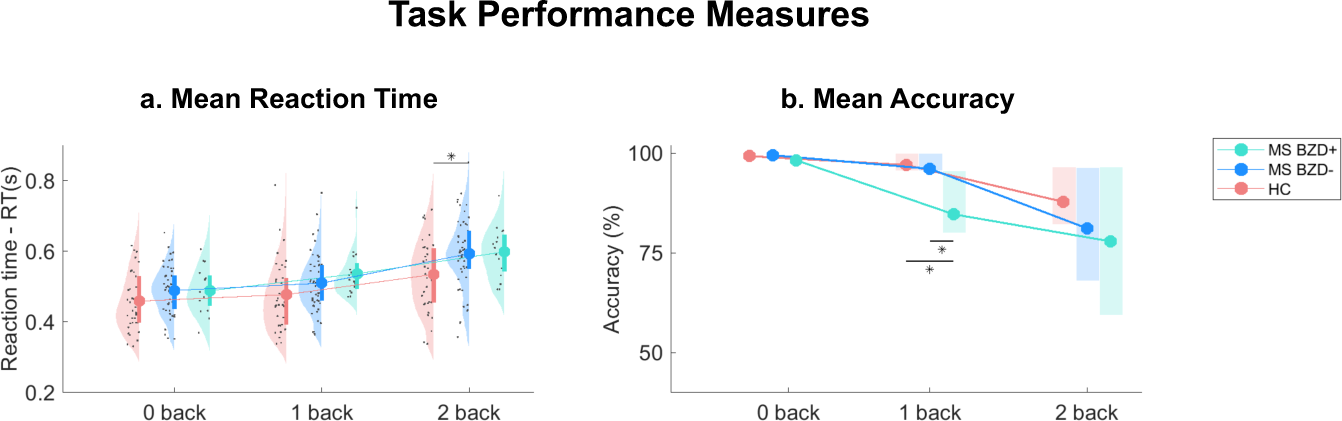
Task Performance measures. A) Reaction time (RT). We plot the distributions of mean RT for the three groups, separately. For each condition, the bold line indicates the inter quartile range [75%-25%]. The dot line tracks the mean for each condition and group. B) Mean Accuracy. The boxes indicate the inter quartile range [75%-25%] of accuracy across subjects for each group. The dot line tracks the mean for each condition and group. In both graphs, the statistics was performed using one-way ANOVA test with the disease condition as grouping variable. The black line between two groups signs the statistical significance, *p value <0.05.

**Figure 4.**
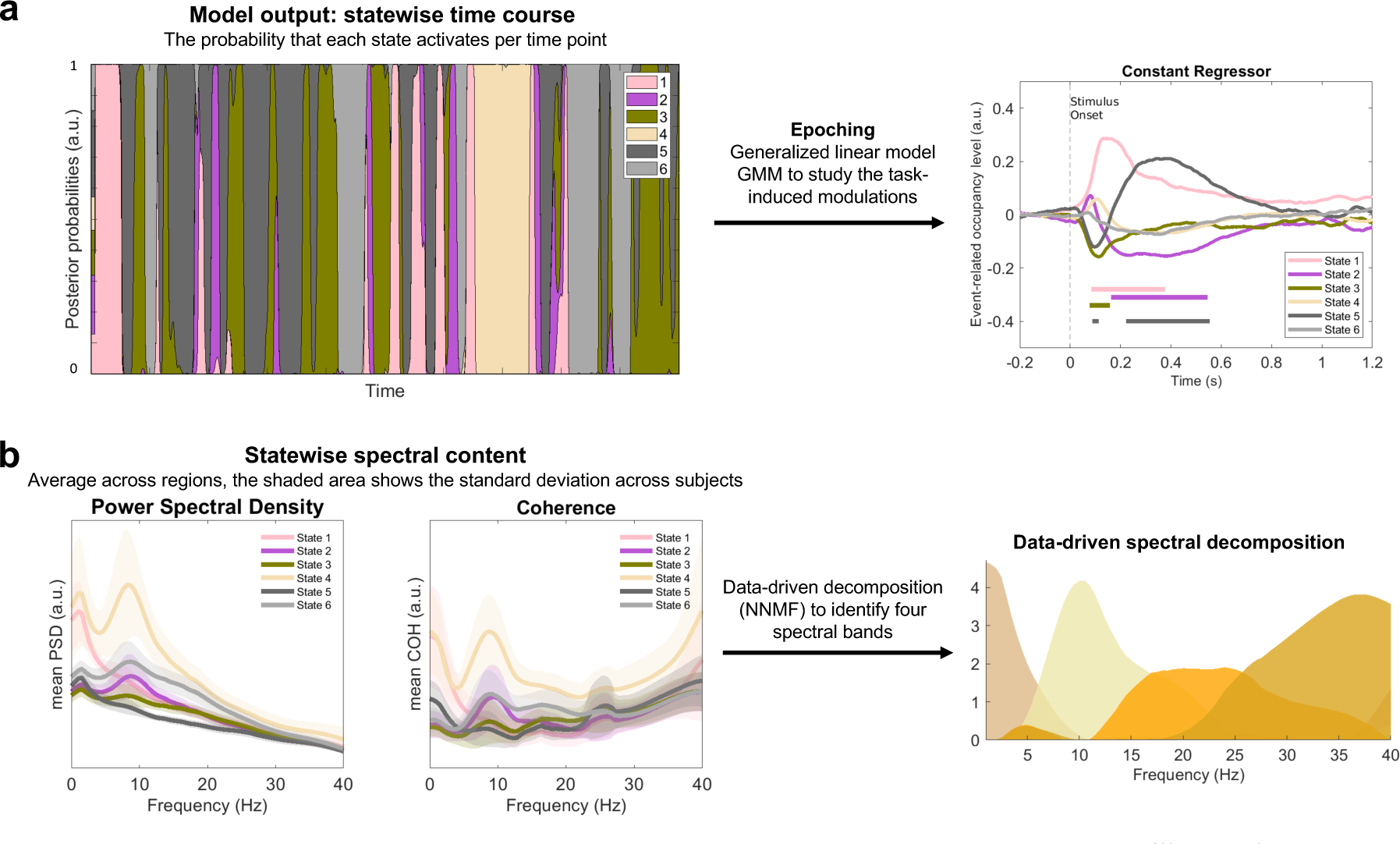
General temporal and spectral descriptions of all the 6 inferred states. a Temporal dimension. We illustrate the steps to extract the event-related activation profiles of all the HMM states. We epoched the statewise time courses with respect to the task information, considering 200 ms before the stimulus onset and 1200 ms after stimulus onset. Then, each trial is baseline corrected considering the [-200, - 30] ms interval as baseline. Finally, we ran the generalized linear model to extract the statewise ER activations. We plotted the constant regressor for all the states, the flat lines underneath the curves represent the time window in which the correspondent state (color-coded) is significantly different from zero, hence, from baseline level. **b Spectral content.** Via multitaper we extracted the statewise power spectral density (PSD) distribution for all regions and the coherence (COH) across all pairs of regions. We plot the average PSD and COH across regions/connections per state (bold line) and the standard deviation across subjects (shaded area). Finally, we decomposed the COH into 4 components via non-negative matrix factorization (NNMF), to identify 4 data-driven frequency bands.

### 3.2 Task-relevant states

To identify the task-relevant states, we plot the constant regressor (the states’ average activity across all paradigm conditions and groups) of the GLM analysis, Figure 4a. We define task-relevant as those states with a modulated

ER activation profile, the ER time course of activation after stimulus onset. We observe that the event-related activation profiles of states 1, 2, 3, and 5 are significantly modulated throughout the epoch (permutation test, number of permutations = 1000, p < 0.025, correction for multiple comparisons via maximum statistics). The modulation is evaluated singularly for each state, and it can be either a positive or negative deflation of the ER curve, denoting respectively a significantly stronger activation or deactivation compared to the baseline level. The ER group comparison is only conducted for the task-relevant states to observe how MS and benzodiazepine treatments affect the WM network dynamics. The description of the non-task-relevant states can be found in Rossi et al.^24^

### 3.3 Spectral modes

Figure 4b illustrates the four data-driven components (spectral modes) in which the broadband frequency range [1-40] Hz was decomposed. We considered these spectral modes as the data-driven frequency bands to refer to when analyzing the states’ spectral content. Spectral mode 1 covers low frequencies (approximately the theta and delta traditional bands 1-8 Hz), spectral mode 2 peaks around 10 Hz (traditionally associated with the alpha rhythm), and spectral mode 3 is broader and covers the beta and low-gamma range (15-40 Hz). Spectral mode 4 then covers mainly higher frequencies (>25 Hz).

### 3.4 Description of the task-relevant states

Following, we present the description of the task-relevant states in the spatial, spectral, and temporal dimensions. The latter regards the statewise event-related (ER) activation profile, reported separately for the three groups (HC, MS BZD-, MS BZD+). Concerning the spectral domain, we identify the spectral mode capturing the state-specific PSD/COH peaks. Finally, for the spectral mode associated with the state, we report the spatial description of the state by meaning of the mean z-score PSD distribution over the brain and phase-coupling network.

#### State 1 – prefrontal theta state

The spectral content of state 1 focuses on the low frequencies (Figure 4b), hence, we associated it with spectral mode 1 covering the traditional delta and theta frequency bands. This low-frequency activity is localized in prefrontal regions (right and left orbitofrontal cortex, OFC, medial prefrontal cortex), the left and right anterior temporal cortices, and the posterior cingulate cortex, PCC, as shown in the PSD map, Figure 5a. The phase-coupling network shows prefrontal-temporal and OFC-PCC connectivity.

**Figure 5.**
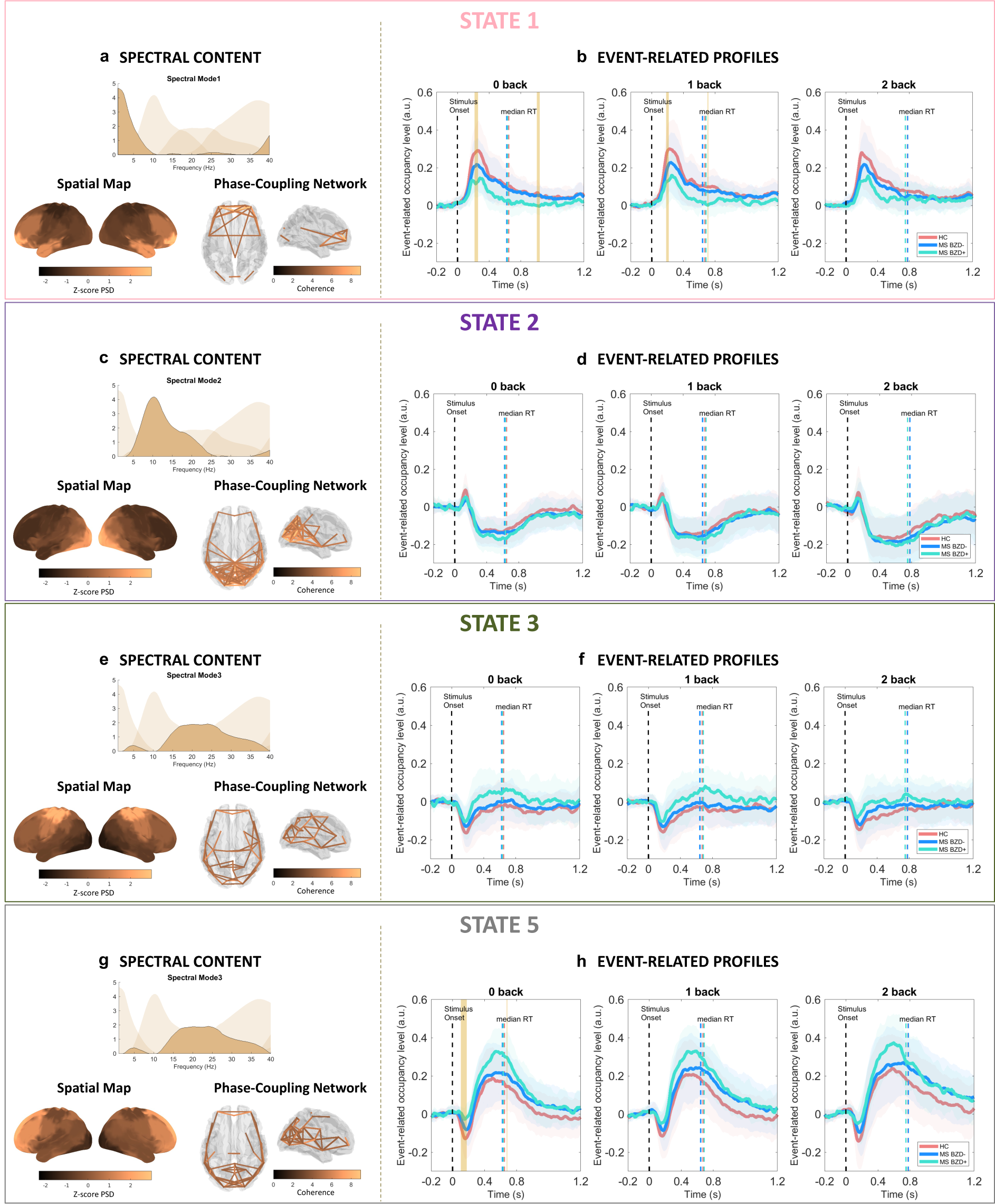
**States Description**. Each box includes the spectral and temporal descriptions of the task-relevant states 1, 2, 3, and 5. Each box presents the spectral content of the states in panels a (for state 1), c (for state 2), e (for state 3), g (for state 5). Each panel presenting the spectral content shows the spectral mode associated with the state, the normalized (z-score) power spectral density (PSD) map, and the brain glass with the phase-coupling network with the connections surviving thresholding via GMM. Then, for each state we report the event-related (ER) profiles in panels b (for state 1), d (for state 2), f (for state 3), h (for state 5). The event-related profiles of activation of each state are reported for the three WM load conditions (0, 1, and 2 back). In each plot, the three ER profiles are color-coded for the three groups (HC, MS BZD-, MS BZD+) and show the mean curve (bold line) and standard deviation (color-coded shaded areas) across subjects. The orange areas delineate the time window in which the three ER waves are significantly different (one-way ANOVA test p<.05, permutation analysis, number of permutations = 1000 and multiple comparison correction via maximum statistics).

This theta prefrontal state is significantly activated early after stimulus onset, peaking ca 180 ms PST (post-stimulus time), Figure 5b. The event-related profiles of this state for the three groups (HC, MS BZD-, and MS BZD+) differ significantly around the maximum peak (one-way ANOVA test, p<.05, number of permutations n = 1000, multiple comparison correction via maximum statistics) in the 0 and 1 back conditions, whilst no significant difference survived in the 2-back condition. The lower ER amplitude entails a decrease in theta coupling between the recruited regions induced by MS and the benzodiazepine treatment.

#### State 5 – M300 frontoparietal state

State 5 presents higher broad spectral activity, with a peak of average COH around 25 Hz, associated with spectral mode 3, capturing the traditional beta band, Figure 4b. This activity is distributed over a frontoparietal network including the inferior and superior dorsal PFC, the medial PFC, the left and right superior PFC, the right and left medial sensorimotor cortex (SMC), and the anterior and posterior precuneus. The synchronization of these regions via the beta rhythm results in the phase-coupling network of state 5, Figure 5g.

The event-related activation of state 5 presents significant modulation across groups, Figure 5h. First, an early negative deflation of the ER wave occurs around 100 ms, and this peak appears less negative for the MS populations; the negative peaks are significantly different (one-way ANOVA test, p<.05, number of permutations n = 1000, multiple comparison correction via maximum statistics) in the 0 back. Following, the event-related wave shows a positive amplitude increase between 300 and 600 ms, which we refer to as the M300 wave as it resembles the typical EEG P300 wave but is extracted in MEG data. This amplitude is significantly different between groups in a short time window around 500 ms in the 0 back condition (one-way ANOVA test, p<.05, number of permutations n = 1000, multiple comparison correction via maximum statistics).

#### State 2 – alpha occipital ac1vity

The spectral content of state 2, both power and coherence, peaks around 10 Hz, associated with spectral mode 2 – the alpha band, Figure 4. This activity is focused in the occipital cortex, and the phase-coupling networks reveal an alpha synchronization between occipital, posterior temporal, and inferior parietal regions, Figure 5c.

The event-related response of this state shows an early non-significant activation around 100 ms post-stimulus presentation, followed by a below-baseline reduction of activity – i.e. deactivation – between 200 and 500 ms, Figure 5d. However, the ER activation profile of this state in the MS group does not deviate from the HC response. Therefore, the activity of this state seems to be affected neither by MS nor BZD administration.

#### State 3 – sensorimotor state

The spectral activity of state 3 peaks around 20 Hz, associated with spectral mode 3, Figure 4b. The regions recruited in this state are the sensorimotor cortices, Figure 5e. The event-related response of this state shows a negative peak around 100 ms post-stimulus, Figure 5f. Qualitatively, we observe that the MS group undergoing benzodiazepine treatment shows a less negative peak that goes back to baseline level faster compared to the HC and MS BZD- groups; however, these differences result statistically non-significant.

### 3.5 Peak analysis

In the peak analysis, we extract the maximum amplitude (activation level) of the statewise ER activation profile and the latency (timing of activation) of the peak. We pursued a peak analysis for the states displaying a significant difference between MS and HC groups. Tables 2, 3, and 4 report the results of the n-way ANOVA test used to investigate the difference between peak amplitudes and latencies between groups and paradigm conditions. All the n-way ANOVA tests are corrected for multiple comparisons via false discovery rate, and we also report the results of Tukey’s multiple comparison tests for the significant ANOVAs.

**Table 2.**
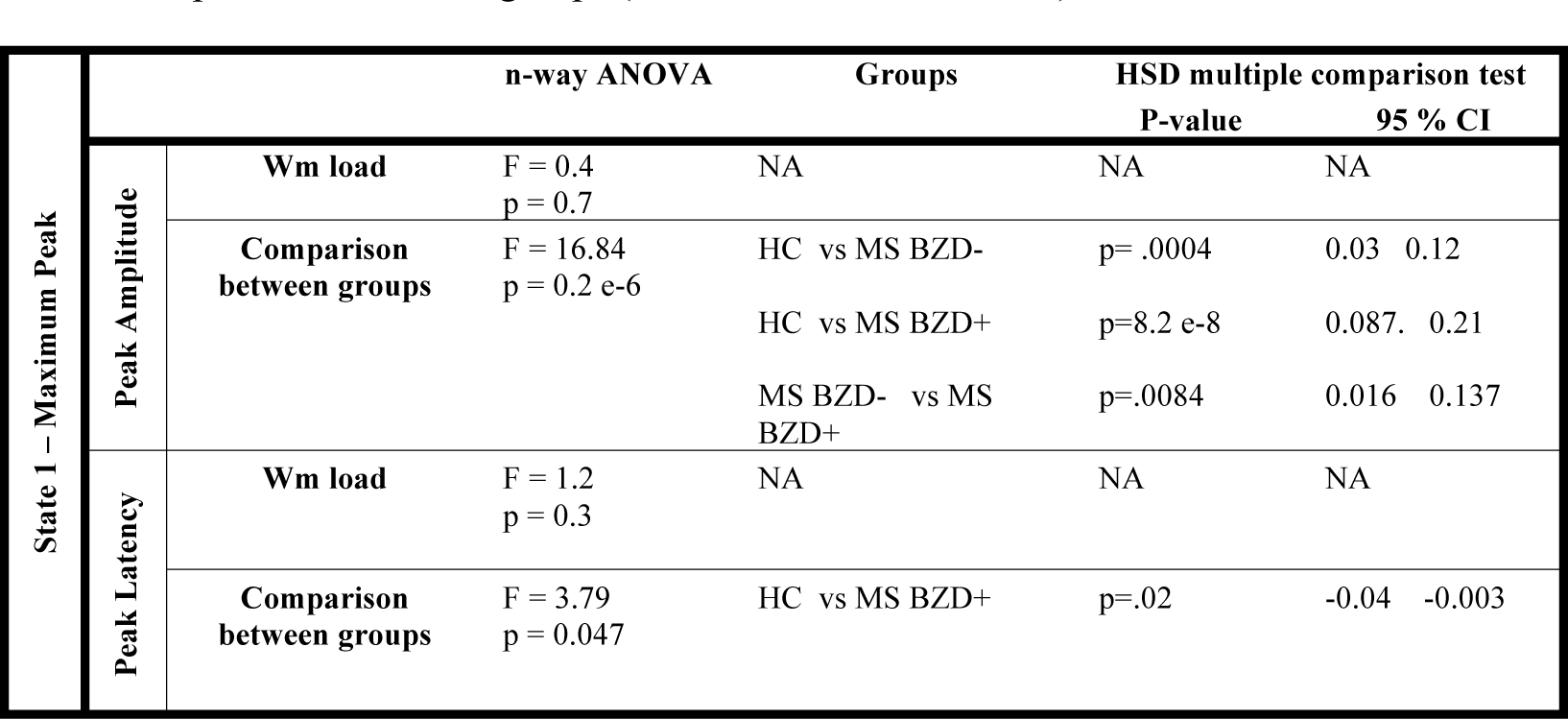
Results of the statistical analysis for the maximum peak of state 1. The table reports the results of the n-way ANOVA test considering 3 groups (HC, MS BZD+ and MS BZD-) and 3 WM load conditions (0, 1, and 2 back). The p-values are corrected for multiple comparison via FDR. For the variables in which the ANOVA test resulted significant, we conducted a multiple comparison analysis via Tukey’s honestly significant difference (HSD) test to observe which comparison between groups lead the difference.

**Table 3.**
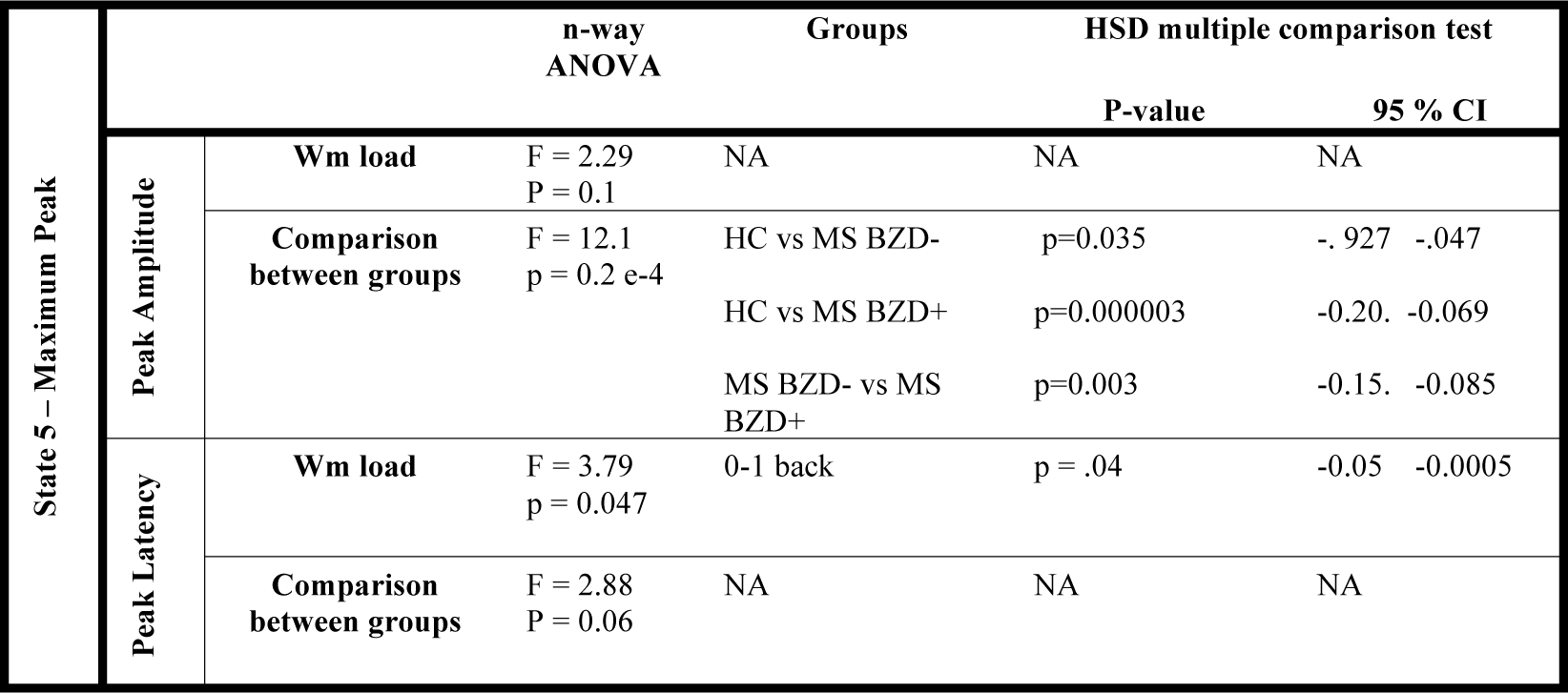
Results of the statistical analysis for the maximum peak of state 5. The table reports the results of the n-way ANOVA test considering 3 groups (HC, MS BZD+ and MS BZD-) and 3 WM load conditions (0, 1, and 2 back). The p-values are corrected for multiple comparison via FDR. For the variables in which the ANOVA test resulted significant, we conducted a multiple comparison analysis via Tukey’s honestly significant difference (HSD) test to observe which comparison between groups lead the difference.

**Table 4.** Results of the statistical analysis for the minimum peak of state 5. The table reports the results of the n-way ANOVA test considering 3 groups (HC, MS BZD+ and MS BZD-) and 3 WM load conditions (0, 1, and 2 back). The p-values are corrected for multiple comparison via FDR. For the variables in which the ANOVA test resulted significant, we conducted a multiple comparison analysis via Tukey’s honestly significant difference (HSD) test to observe which comparison between groups lead the difference.

|  |  | n-way ANOVA | Groups | HSD multiple comparison test |  |
| --- | --- | --- | --- | --- | --- |
|  |  |  |  | P-value | 95 % CI |
| Peak Amplitude | Wm load | F = 1.89<br>P = 0.15 | NA | NA | NA |
|  | Comparison between groups | F = 14.97<br>P = 0.12 e-5 | HC vs MS BZD- | p = .001 | -0.07 -0.014 |
|  |  |  | HC vs MS BZD+ | p = .5 e-6 | -0.12 -0.046 |
|  |  |  | MS BZD- vs MS BZD+ | p = .01 | -0.078 -0.0073 |
| Peak latency | Wm load | F = 0.45<br>P = 0.6 | NA | NA | NA |
|  | Comparison between groups | F = 4.93<br>P = 0.02 | MS BZD- vs MS BZD+ | p = .005 | 0.0055 0.0382 |

The comparison between WM load conditions is significant only for the latency of the maximum peak of state 5. Instead, we observe that the comparisons between groups (HC, MS BZD-, MS BZD+) show significant amplitude and latency differences for all the peaks, both for states 1 and 5. Figure 6 plots the distributions of the amplitudes and latencies of all peaks for the three groups (HC, MS BZD-, MS BZD+).

**Figure 6.**
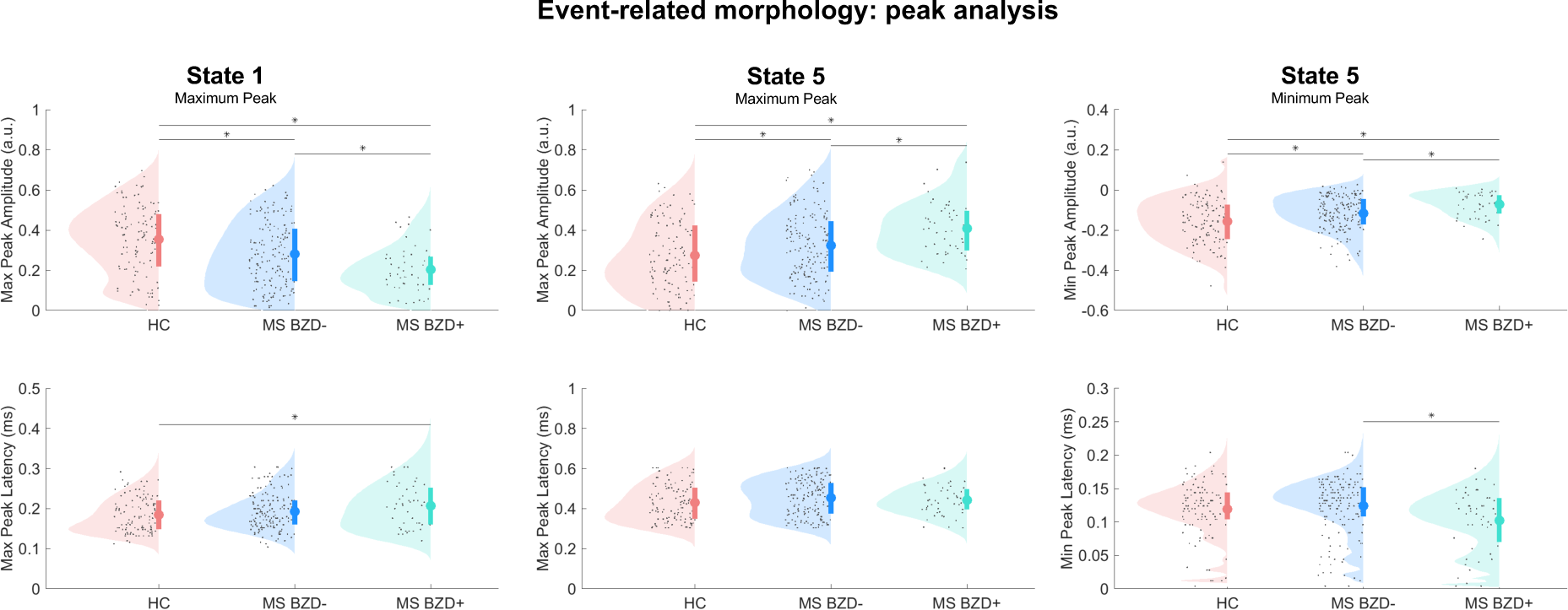
Distributions of the peaks’ amplitudes and latencies for the three groups HC, MS BZD-, and MS BZD+. Each plot presents the distributions of peak amplitude (top row) and peak latencies (bottom row) for the three groups under analysis for the maximum peak of state 1 (a), the maximum peak of state 5 (b) and the minimum peak of state 5 (c). The significance level is drawn with a line between the two groups that differ. *p<.05 (for the precise p value we refer to Tables 2, 3, and 4). The significance levels are computed via n-way ANOVA with multiple comparison correction via false discovery rate, and the post-hoc analysis via Tukey’s multiple comparison test.

### 3.6 Relationship between states’ ER significant features and behavioral data

Figure 7 reports the correlations between the behavioral data and the ER features of state 1, and Figure 8 reports the correlations between the behavioral data and the ER features of the M300 peak for state 5. The maximum and minimum peaks of state 5 display the same behavior, thus, we only considered the maximum peak of state 5. For this analysis, we only considered the MS BZD- group to discard any BZD-induced effect and observe whether the ER features capture MS-related aspects of the brain dynamics directly linked to behavioral data.

**Figure 7.**
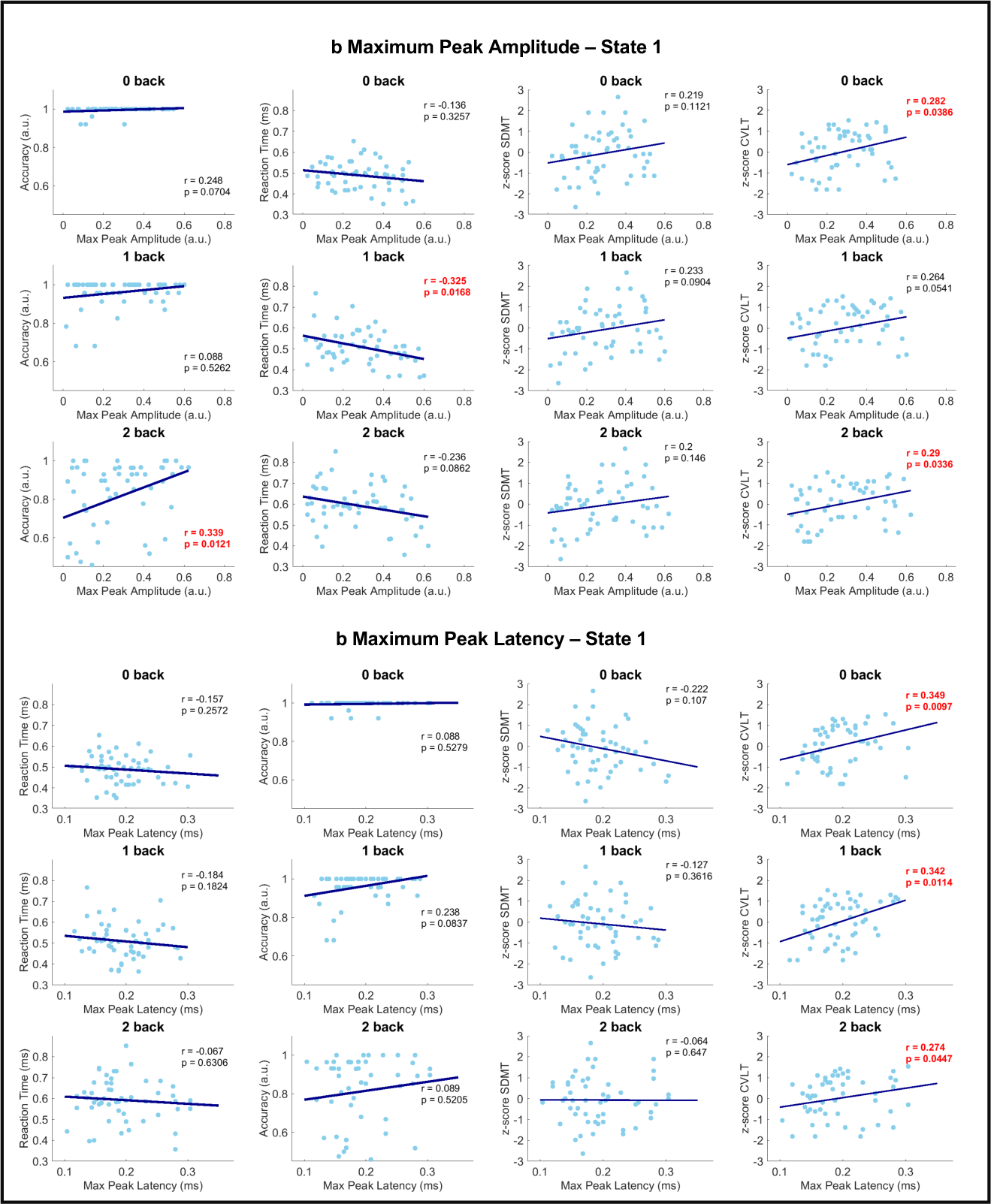
Relationship between state 1 ER peaks and behavioral data for the MS cohort without benzodiazepine treatment. The behavioral data include the mean reaction time and accuracy per subject, and two neuropsychological tests, the single digit modality test (SDMT) and the California Verbal Learning Test-II (CVLT-II). The scores of the neuropsychological tests are standardized (z-score). a Correlation between the amplitude of state 1 maximum peak and the behavioral data. b Correlations between the latency of state 1 maximum peak and the behavioral data. All the correlations are computed as Pearson’s correlations, and the p-values are not corrected for multiple comparisons. In each plot, the correlation coefficient (r) and the p_values (p) are displayed. Each row presents the data for a single WM load condition.

**Figure 8.**
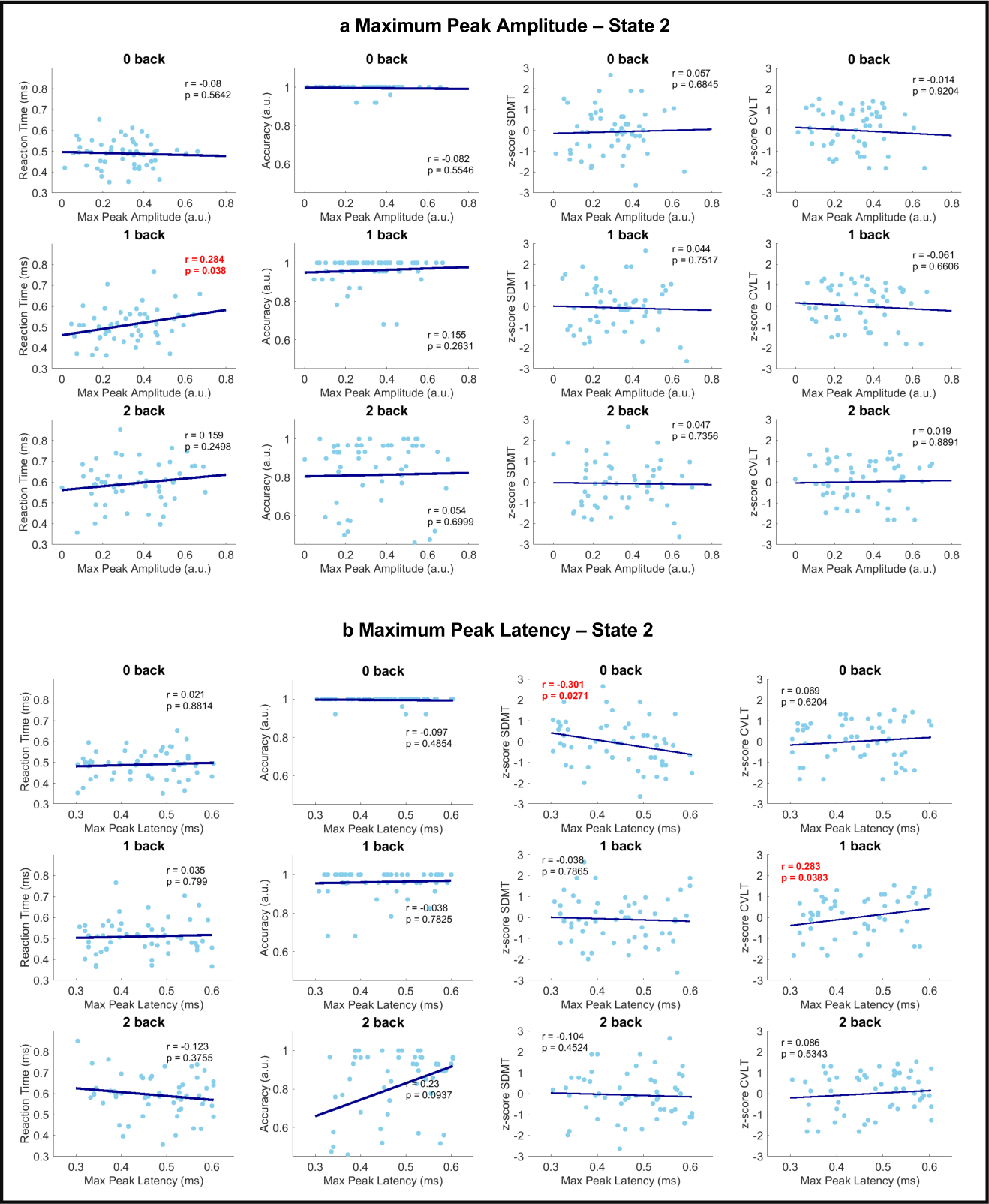
Relationship between state 5 ER peaks and behavioral data for the MS cohort without benzodiazepine treatment. The behavioral data include the mean reaction time and accuracy per subject, and two neuropsychological tests, the single digit modality test (SDMT) and the California Verbal Learning Test-II (CVLT-II). The scores of the neuropsychological tests are standardized (z-score). a Correlation between the amplitude of state 5 maximum peak and the behavioral data. b Correlations between the latency of state 5 maximum peak and the behavioral data. All the correlations are computed as Pearson’s correlations, and the p-values are not corrected for multiple comparisons. In each plot, the correlation coefficient (r) and the p_values (p) are displayed. Each row presents the data for a single WM load condition.

For state 1, we observe that both peak amplitudes and latencies are positively correlated with the CVLT scores in all paradigm conditions (except for the 1 back for the amplitude). Additionally, the peak amplitude of state 1 is significantly negatively correlated with the mean reaction time in the 1 back and significantly positively correlated with the mean accuracy in the 2-back condition. Instead, for state 5, we report a significant negative correlation between the peak latency and the SDMT score in the 0-back condition. In this exploratory analysis, we did not apply correction for multiple comparisons.

## 4. Discussion

Through the time delay embedded-hidden Markov model (TDE-HMM), we decomposed the task MEG data in spectrally-defined and data-driven functional brain networks.^25,30^ We previously implemented this technique to unveil the WM network dynamics in a healthy population, and we identified four task-relevant brain networks.^24^ Expanding the analysis to the MS cohort, we observed that the disease itself and the ongoing benzodiazepine treatment affect the dynamics of two networks: the theta prefrontal and the M300 frontoparietal networks.

The spatial configurations of states 1 and 5 include frontal, temporal, and parietal regions, which majorly contribute to normal WM functioning.^7^ FMRI studies have consistently reported an MS-induced altered activation of the same regions during WM tasks.^5,10^ The link between the HMM states, the WM networks, and MS findings supports the model reliability in detecting functional networks relevant to investigating WM and disease-specific processes.

However, the fMRI literature reports contradictory findings on the altered network activation.^5,37^ Whilst the increased activity of the prefrontal cortex in pwMS has been explained as an effect of functional reorganization to compensate for structural damage^5,8,38^, a decreased frontoparietal activation and coupling were found to correlate with decreased WM task performance.^9,39,40^ These discrepancies might arise from the heterogeneity of the datasets, as Vacchi et al.^10^ demonstrated that the pattern of altered network activation strongly depends on the MS phenotype (relapsing-remitting or progressive) and disease stage.^10,39^ Therefore, in our research, we confined the analysis to relapsing-remitting pwMS and further split the MS cohort in pwMS treated with (MS BZD+) and without (MS BZD-) benzodiazepines (BZDs), widely used symptomatic treatments in MS care to reduce anxiety, muscle spasticity, and insomnia. Although the neurophysiological alterations of benzodiazepine have previously been observed in HC^27,41^, MS studies have never considered this treatment as a separate contrast, which might have also led to discrepant findings.^20^ We start by discussing the dynamics of the MS BZD- group, elaborating on the BZD effect separately.

### 4.1 State 1 – impaired theta prefrontal network

State 1 depicts the early low-frequency (theta) prefrontal activity. Generated by the hippocampal-cortical circuit, theta is an excitatory rhythm that supports high-order cognitive functions, such as information integration between executive (prefrontal) and maintenance (temporal) regions, and the top-down attentional control from low to high cognitive processing (the OFC-PCC theta coupling).^42–44^ Additionally, state 1 peaks at about 180 ms after stimulus onset, suggesting that the state is activated in the first encoding stage of WM processing.^24^ Therefore, we hypothesize that state 1 performs early executive functions, crucial in (verbal) encoding and maintenance processes during a visual-verbal n-back task.^24^

The significantly decreased activation of state 1 in pwMS suggests a reduced theta coupling between the regions in pwMS compared to HC. These findings are complementary to the more traditional time-frequency analyses, reporting a decreased theta power in the hippocampus and frontal regions of pwMS.^16,45^ The network configuration that that this work describes provides additional information about the underlying functional mechanisms impaired in MS: the decreased theta activity of state 1 indicates a decreased functional (dictated by the theta rhythm) integration between regions, consistently to what previously reported in more static graph theory analyses.^40,46^

The decoupling of state 1 could result from an underlying structural damage due to lesions and atrophy, yielding functional connectivity damages. A longitudinal study reported that MS lesions develop preferentially in frontal and parietal regions, suggesting a link to the early WM impairment affecting pwMS.^47,48^ However, lesion development is extremely subject-dependent. Therefore, future studies should investigate the relationship between altered dynamic functional networks and lesion load and location.

Viewing the neuropsychological multi-compartment model of WM, we could link state 1 to the executive unit, given the state’s encoding and attentional control functions.^6^ The effect of MS on the activation of state 1 corroborates the hypothesis that the executive control unit is more strongly affected by MS than the two slave units, as previously suggested by neuropsychological research.^4,8,49^ Consistently, during the visual-verbal n-back task, the recruited slave unit is the phonological loop captured by state 2 (see ref.^24^), and we observe that its activation does not change in pwMS. This link to neuropsychology is enabled by the use task data. Van Schependom et al. utilized the same TDE-HMM technique on resting-state MEG data and identified a frontal DMN with reduced theta power in MS compared to HC.^20^ As explained in Rossi et al., state 1 presents spatio-spectral features that resemble the frontal default mode network (DMN), in particular, the theta coupling between the orbitofrontal and posterior cingulate cortices.^24,25^ However, performing the analysis on task data allows us to link the HMM state, hence the functional network, with the specific cognitive process required during the task.

The peak analysis reveals that a correctly and rapidly performed task requires a strong activation of state 1, particularly with increasing task difficulty. This result supports the same hypothesis for which WM performance significantly relies on the well-functioning of the executive control unit.^49^ We also observe that the peak amplitude and latency of state 1 correlate with the California verbal learning test-II (CVLT), a neuropsychological test to assess the initial verbal learning process. This suggests that a successful stimulus encoding in MS requires higher ER activation of the theta prefrontal network, which, however, seems to have a delayed latency, probably reflecting the underlying structural damage. These results align with previous neuropsychological findings, suggesting that (working) memory impairment was associated with difficulties in the initial learning rather than the following retrieval stage^50^, as we discuss in state 5.

### 4.2 State 5 – an MEG M300 frontoparietal network

State 5 captures a beta frontoparietal network with an ER temporal activation profile that resembles the widely used and detected EEG P300. This characteristic wave is associated with the stimuli discrimination process and represents a cognitive marker for several high-order cognitive functions, such as attention and working memory.^51^

The ER wave of state 5, the M300, does not seem to be majorly altered in the MS BZD- group compared to the HC. These results contradict the main findings in the MS P300 literature that repeatedly reports decreased P300 amplitudes or increased P300 latencies in pwMS.^15,48,52,53^

The discrepancies that we report may arise from methodological and signal-specific issues. First, the biophysical difference between the electric and magnetic signals causes the MEG and EEG to be sensitive to different neuronal orientations, making the MEG and EEG acquisitions refer to slightly different brain sources.^54^ Secondly, the EEG signal suffers from volume conduction, thus, the electric signal does not homogeneously travel across the different head components (brain, meninges, skull).^54^ As traditional EEG analyses are conducted at the sensor level, the EEG P300 could result from more widespread changes that sum up at the scalp level resulting in a P300 reduction in patients with MS. Instead, source-level event-related activations are more sensitive to region-specific phenomena.^16^ Therefore, although our M300 results seem to clash with the traditional EEG P300 literature, these cannot be straightforwardly compared, as they observe the event-related brain response from different perspectives: source (MEG) versus sensor (EEG) level and network (TDE-HMM) versus single-region (traditional ER) activity.

Considering the relationship with behavioral data, our results suggest that an increased M300 latency indicates a decreased information processing speed (SDMT score), as previous EEG P300 studies have also reported^55^. Therefore, the latency of the M300 state 5 seems to share the same neurophysiological meaning as the EEG P300 latency.

Instead, regarding the amplitude, the M300 shows a weak increased amplitude in pwMS, indicating a more consistently activated beta activity in pwMS than HC. This is not the first time that an event-related analysis has reported an increased beta activity in pwMS, as Kiiski et al reported the same effect in EEG source-reconstructed data.^45^ However, it is difficult to link this finding to an altered MS-related mechanism, and future studies should further investigate this beta-related effect.

### 4.3 Benzodiazepines

Several electrophysiological (M/EEG) studies have previously observed the effect of benzodiazepines on healthy brain activity, both in event-related response and oscillatory dynamics^27,41,55^. BZDs increase the GABAA conductance of inhibitory interneurons^41^, which take part in the circuitry generating the theta and beta oscillatory activity^41–43^. Considering the MS BZD+ group, our results show an altered ER activation profile for states 1 and 5, the networks characterized by theta and beta activity, respectively.

The ER activation of state 1 in the MS BZD+ group is significantly decreased compared to HC and MS BZD-. This alteration entails a further reduced theta power activation in state 1, caused by BZD use. Van Schependom et al. reported the same effect in resting-state MEG data.^20^ The pharmacological effect of benzodiazepine induces an increase in inhibitory currents in the GABAergic inhibitory interneurons, disrupting the excitation-inhibition balance of the cortical (medial septum-diagonal band of Broca)-hippocampal circuitry that generates the theta rhythm.^42,43,56^ Therefore, what we and Van Schependom et al. reported are the macroscopic network alterations reflecting the BZD effect at the neuronal circuitry level.

Instead, the beta rhythm is generated by local circuits of GABAergic inhibitory interneurons in the primary somatosensory and sensorimotor cortices^57^. The increase in beta power derives from the increased GABAergic inhibition, that has systematically been reported in pharmacological, EEG, and MEG studies^27,41,56,58^. Van Schependom et al. found a BZD-induced increased beta power in several TDE-HMM states^20^. Our results show the same reoccurring effect in the two states characterized by beta activity, states 3 (only qualitatively) and 5. This last depicts frontoparietal beta activity and presents a significantly increased M300 activation in the MS BZD+ group compared to HC and MS BZD-.

Our results on the M300 clash with the EEG P300 findings in the BZD literature. Many studies report a decreased P300 amplitude as the effect of benzodiazepine^27,55,59^. These discrepancies might arise from the same methodological issues that underlie the discrepant M300 findings between MS BZD- and HC. To the best of our knowledge, there is no understanding of the neural mechanism that generates the BZD-induced decrease in P300 amplitude. Instead, our approach enables us to link the phenomenon that generates the M300 wave and the underlying neuronal mechanism of its alteration.

Concerning the behavioral effects, BZD use was associated with reduced accuracy in performance in HC^55,58^. In our results, the accuracy in the task performance is significantly altered in the MS BZD+ group compared to the MS BZD- group, Figure 3b. Both states 1 and 5 carry out executive functions that can impact the task performance if altered, which becomes the case in patients with MS undergoing benzodiazepine treatment. We hypothesize that the altered benzodiazepine-induced activation observed in our results plays a role in performing the task accurately. However, due to the limited sample size (only 16 pwMS with benzodiazepine) we couldn’t further investigate the relationship between BZD use, TDE-HMM states 1 and 5 ER activation profiles, and task performance measures.

## Limitations and future work

We explored the statewise event-related response that provides the first level of information on connectivity and power changes for each network. As we inferred the HMM states on the concatenated data (HC and MS), the model inferred the states at the group level. This choice allowed a direct comparison between event-related responses across groups, obscuring connectivity changes between groups. If we had inferred HMM states separately for the different cohorts, we would need to run an additional step to identify which state in one inference relates to which state in the other inference. Due to the stochastic nature of the HMM inference, chances are that the states between inferences slightly differ, hindering the investigation of the disease-induced alterations. Future works should evaluate a way to pursue a more detailed connectivity analysis between groups.

We did not include structural data such as lesion load, atrophy measures, or information about the lesion locations. Therefore, we only speculated on the link between the altered activation of state 1 and the structural measures, and future studies should investigate this relationship at the group and subject levels. Further studies should also explore the variation in brain response between different benzodiazepines.

## Conclusions

Our study examined how, during a visual-verbal WM n-back task, MS and BZDs treatment affect the event-related response of spectrally defined functional networks identified through the unsupervised TDE-HMM approach. We found that MS impairs the activation of a theta prefrontal network associated with early stimulus encoding and attentional control. This result resonates with the neuropsychological literature explaining WM impairment in MS as a difficulty in the early verbal learning of WM processing.

Treatment with BZDs caused a further decreased activation of the theta prefrontal network and an increased activation of beta frontoparietal M300 network. These macroscopic alterations of the network dynamics reflect the treatment effect on the neuronal circuitries generating the theta and beta rhythms, respectively. Our study demonstrates that the model can identify and differentiate the disease-specific and treatment-specific effects, revealing potential new markers to assess the condition of working memory in multiple sclerosis.

## Data availability

The MEG and MRI data for this study are not publicly available due to privacy restrictions. Researchers interested in collaborating on these data are welcome to contact the senior authors (Prof. Jeroen Van Schependom and Prof. Guy Nagels).

## Code availability

The analyses were conducted in MATLAB, utilizing the freely accessible HMM-MAR package which can be found here: https://github.com/OHBA-analysis/HMM-MAR. This package belongs to the OSL (OHBA Software Library) toolbox that can be consulted here: https://ohba-analysis.github.io/osl-docs/. In particular, the analysis we implemented was based on the work presented by30. The scripts containing the full pipeline (MEG preprocessing, HMM inference, and data analysis with GLM and spectral decomposition), and that can be used to reproduce the analysis conducted in this work, can be found here: https://github.com/OHBA-analysis/Quinn2018_TaskHMM. For more details on the analysis scripts, contact the corresponding author (Chiara Rossi).

### Acknowledgements

The authors would like to thank the participants for their time and commitment to this study. The MEG data collection was enabled by grants from the Belgian Charcot Foundation and by an unrestricted research grant provided by Genzyme-Sanofi. C.R. is funded by Fonds Wetenschappelijk Onderzoek (FWO, Grant numbers: 11K2823N, 11K2821N). Author contributions C.R. conducted the analysis and wrote the manuscript. J.V.S. was the main supervisor of the work and helped write and review the manuscript. D.V., L.C., and G.N. gave inputs for the analysis and provided feedback in the writing process. M.W., M.B.D., M.D and F.A. provided comments on the work. All authors approved the submitted version.

